# Iron-responsive phosphorylation of TolQ modulates cell envelope integrity and antibiotic susceptibility in *Klebsiella pneumoniae*

**DOI:** 10.64898/2026.04.25.720785

**Authors:** Chelsea Reitzel, Jonathan Sayewich, Stevan Cucic, Oscar Romero, Norris Chan, Jennifer Geddes-McAlister

## Abstract

*Klebsiella pneumoniae* is an opportunistic bacterial pathogen associated with high morbidity and mortality, exacerbated by the rapid emergence of resistance to last-resort antibiotics, such as carbapenems. Adaptation to nutrient limitation, particularly fluctuations in metal availability, is critical for bacterial survival and virulence, yet the regulatory mechanisms coordinating these responses remain incompletely understood. Protein phosphorylation represents a key post-translational modification governing bacterial physiology and offers a promising avenue for identifying novel antimicrobial targets. Here, we applied mass spectrometry–based phosphoproteomics to define nutrient-responsive signaling networks in *K. pneumoniae* under varying iron and zinc conditions. This analysis identified iron-dependent phosphorylation of TolQ, a conserved inner membrane component of the Tol–Pal system that maintains cell envelope integrity. Structural modeling predicted that phosphorylation modulates TolQ–TolR conformation, suggesting a mechanism by which iron availability regulates Tol–Pal function. Functional characterization demonstrated that deletion of *tolQ* results in reduced bacterial viability, increased susceptibility to host immune clearance, and heightened sensitivity to antibiotic treatment. To further explore the therapeutic potential of this pathway, we integrated high-throughput compound screening with computational modeling and identified small molecules that phenocopy Δ*tolQ*. Collectively, these findings reveal a previously unrecognized link between iron availability and phosphoregulation of the Tol–Pal system and establish TolQ as a critical mediator of bacterial survival. This work highlights phosphoproteomics as a powerful strategy to uncover regulatory vulnerabilities and identify targets for antimicrobial development in drug-resistant pathogens.

## Introduction

*Klebsiella pneumoniae* is a human bacterial pathogen often associated with hospital-acquired infections causing pneumonia and bacteremia, leading to mortality rates reaching 20% (1). Critically, *K. pneumoniae* has developed resistance to antibiotics, such as β-lactams, including carbapenems, which are designated as “last resort” antibiotics, limiting therapeutic options and doubling mortality rates in the wake of these ineffective therapies (2). A recent meta-analysis of global *K. pneumoniae* infections showed that 28% of strains were carbapenem-resistant, with several geographical regions finding resistance in 70% of strains (3). The health implications of disease and rising rates of resistance underscore a need for novel antimicrobials or strategies for resensitization of the pathogen for improved global health. One such approach is the targeting of proteins that mediate bacterial adaptation to nutrient availability. For example, proteins regulated by iron and zinc are particularly compelling candidates, given the critical roles of these metals in microbial growth, cell division, biofilm formation, and virulence (4–12), processes previously targeted by existing antibacterial agents (13, 14). To survive in nutrient-limited environments, bacteria have evolved complex mechanisms, including phospho-signaling, to adapt to fluctuations in nutrient supply (15). Characterizing the phosphorylation profile of *K. pneumoniae* under varying iron and zinc conditions can uncover regulatory nodes and protein targets that can be exploited as novel antimicrobial interventions.

Protein phosphorylation is a reversible post-translational modification that alters protein activity, stability, or subcellular location through the addition of negatively charged phosphate groups. In bacteria, phosphorylation commonly occurs on serine, threonine, tyrosine, aspartic acid and histidine residues, and more rarely, cysteine (16), arginine (17), lysine (18) and glutamic acid (19). This modification regulates many physiological processes in bacteria, including processes involved in infection and colonization of a host (20, 21). Notably, the mechanisms of phosphorylation in bacteria differ substantially from those in eukaryotes, making bacterial kinases and their substrates attractive targets for novel antimicrobial strategies (22). Consequently, phosphoproteomic profiling provides a powerful approach to understanding bacterial virulence and identifying potential therapeutic targets. Moreover, advances in mass spectrometry coupled with enrichment techniques, such as immobilized metal affinity chromatography (IMAC), have enabled high-throughput identification and quantification of phosphoproteins. Numerous bacterial phosphoproteome datasets have been reported (23–28); however, to date, there has not been a proteome-wide profiling on the effect of iron and zinc availability on phosphoproteome in *K. pneumoniae*, which would provide insight into regulatory roles of the modification with potential implications for discovery of bacteria-specific therapeutic targets.

Within the present study, our investigation of phosphoregulation of *K. pneumoniae* upon modulation of nutrient availability uncovered a role for TolQ, a conserved inner membrane protein in Gram-negative bacteria, involved in maintaining membrane integrity. Specifically, TolQ belongs to the Tol-Pal system, a multi-protein complex (TolA, TolB, TolQ, TolR, Pal) that stabilizes the bacterial cell envelope (29). TolQ forms a pentameric pore and, in complex with TolR, acts as a proton-driven motor to enable interactions between TolA and the TolB-Pal complex (30). TolA facilitates the release of TolB from Pal, regulating Pal mobilization (31). Pal, anchored in the outer membrane via an N terminal lipid moiety, accumulates at division sites where its periplasmic domain establishes noncovalent links to the peptidoglycan layer, supporting the outer membrane during constriction (31, 32). Disruption of Tol genes in *Escherichia coli* leads to membrane leakage and antibiotic sensitivity (33). The system also serves as a receptor for bacteriophages and colicins (34), highlighting its dual role in bacterial membrane integrity and vulnerability. Herein, we present a previously unreported connection between iron availability and phosphoregulation of TolQ and functional assessment of the deletion strain in *K. pneumoniae*. We demonstrate through predictive modeling that TolQ-TolR conformation is modulated by TolQ phosphorylation, which occurs exclusively in iron-replete media, suggesting a regulatory link between iron, phosphorylation, and TolQ function. Phenotypic characterization revealed that Δ*tolQ* exhibited significantly reduced viability, heightened susceptibility to host immune clearance, and increased sensitivity to existing antibiotics. Furthermore, high-throughput compound screening integrated with deep learning, identified small molecules that phenocopy Δ*tolQ*, highlighting their potential as novel antibiotic candidates. These findings underscore the power of phosphoproteomics integrated with phenotypic profiling, high-throughput screening, and computational predictive modeling to uncover targets critical for bacterial survival and pathogenesis, accelerating discovery of targeted antimicrobial therapies against clinically relevant pathogens.

## Materials and Methods

### Bacterial strains and media preparation

*Klebsiella pneumoniae* wild type (WT) K52 serotype (Kp52.145) was maintained on Lysogeny broth (LB) agar plates and Δ*tolQ* was maintained on LB with 34 µg/mL chloramphenicol.

M9 minimal media (Difco TM M9 salts, 0.4% glucose, 2 mM MgSO_4_, 0.1 mM CaCl_2_, chelex MQ H_2_O) and low salt LB media (5% yeast extract, 10% tryptone, 0.5% NaCl, 0.5 µM EDTA) were used as indicated throughout. Preparation of strains, cultures, and *in vitro* experimentation performed as described (35), with study-specific modifications reported.

### Sample preparation for LC-MS/MS analysis

Overnight cultures in 5 mL LB medium were prepared in quadruplicate and incubated at 37 °C. Cells were harvested by centrifugation at 3,500 × g for 10 min and washed twice with 1 mL M9 minimal media and resuspended in 1 mL M9 minimal media followed by subculture at a 1:100 dilution into 100 mL of: M9 minimal media (LM), M9 minimal media supplemented with 10 µM iron (Fe_2_(SO_4_)_3_; iron-replete) and M9 minimal media supplemented with 10 µM zinc (ZnSO_4_; zinc-replete). Cultures were incubated for 24 h at 37 °C to reach late stationary phase and collected by centrifugation. The cell pellets were washed with phosphate-buffered saline (PBS) and processed as previously described (36, 37). The pellets were resuspended in 100 mM Tris-HCl (pH 8.5) with 2% sodium dodecyl sulfate, and protease inhibitor (Roche) and PhosSTOP (Roche) tablets. Cells were lysed by probe sonication, followed by reduction (10 mM dithiothreitol) and alkylation (55 mM iodoacetamide), and precipitation by chloroform–methanol extraction. Phase separation used 0.6 mL Milli-Q water (the upper aqueous phase was removed) with methanol added, vortexing, and centrifugation at 10,000 × g for 5 min to pellet proteins. The pellet was washed (0.4 mL methanol), dried at 45 °C for 10 min, and resuspended in 8 M urea/40 mM HEPES buffer for quantification BSA–tryptophan assay (38). Samples were digested with trypsin/LysC 1:50 enzyme-to-protein ratio (room temperature, overnight) and desalted using peptide desalting columns (Thermo Fisher Scientific, 89852) according to the manufacturer’s instructions. Peptides were divided, with 10% (∼100 µg) reserved for total proteome analysis and dried to completion, while the remaining 90% (∼900 µg) was used for phosphopeptide enrichment.

### Phosphopeptide enrichment

Phosphopeptides were enriched as previously described with minor modifications (36, 37, 39). Peptides were reconstituted in 0.2 mL Binding/Wash buffer (Thermo Fisher Scientific, A32992), adjusted to pH 2.5, and incubated in ferric nitrilotriacetate (Fe-NTA) columns (equilibrated with Binding/Wash buffer) for 15 min (40). Columns were washed (with Binding/Wash buffer and Milli-Q water) and phosphopeptides were eluted using 0.2 mL Elution buffer and dried completely in a SpeedVac at 45 °C.

### LC/MS-MS

Total proteome (5 µg) and phosphoenriched (90% sample volume) samples were reconstituted in 40 µL 0.1% formic acid (FA) and loaded onto Evotips according to the manufacturer’s instructions (41). Samples were analyzed using Themo Scientific Orbitrap Exploris 240 coupled with Evosep One liquid chromatography system using a 44 min (phosphoenriched samples) and 88 min (total proteome) gradient. The mass spectrometer was operated in data-dependent, positive ion mode using a 15 cm PepSep column with a 150 µm diameter and 1.9 µM beads (Evosep, EV1106). The precursor range (350-1800 *m/z*) with a resolution of 60,000, 50 msec injection time, and an automatic gain control (AGC) target of 3e6 were set. Dynamic exclusion (10 sec) and charge states (2–8) were included. The fragment ion isolation window used higher-energy collisional dissociation (HCD) fragmentation energy of 34 eV with a 1 *m/z* window and resolution (15,000) and dynamic injection time.

### Raw data processing

MaxQuant (v2.4.0.0) (42) with default parameters except where noted was used for processing of .RAW mass spectrometry files. Peptide identification was performed using the built-in Andromeda (43) search engine with *K. pneumoniae subsp. pneumoniae* K52 serotype proteome (5126 protein sequences from UP000000265 proteome ID; accessed Dec. 2, 2022) from UniProt. Variable modifications included: phosphorylation of STY, pHis and pAsp modifications (neutral loss of H_3_O_4_P; mass 97.9768950 Da) and a diagnostic peak for pHis (C_5_H_8_O_3_N_3_P immonium ion with a mass of 190.037604 *m/z* after protonation), methionine oxidation, and N-acetylation. Fixed modifications included: carbamidomethylation of cysteine. Abundance of phosphopeptides was normalized to the total proteome with modified and unmodified peptides used for protein quantification by label-free quantification (LFQ) (ratio count of 1) (44). Digestion parameters: trypsin specificity with a maximum of 2 missed cleavages; minimum number of peptides: 2; and “match between runs”: enabled (match window time of 0.7 min and alignment time window of 20 min) (44).

### Bioinformatics

Phospho (STYDH) modifications were processed with Perseus (v2.0.10.0) (45) incorporating phosphorylation multiplicities of 1, 2, and 3, with peptides matching contaminants or reverse peptides removed. Phosphopeptides were classified as: class I (localization probability >75%), class II (>50%), and class III (<50%). Valid value filtering of 3 values in at least one group was set for phosphopeptide LFQ intensities, missing values were imputed by Gaussian distribution (standard deviation width of 0.3 and a downshift of 1.8), and statistical testing by a Student’s *t*-test for false discovery rate (FDR) corrected p-values (at 5%) and S_0_ = 1, were considered significant. Visualizations were performed using Perseus (45) and ProteoPlotter (46).

### Generating competent K. pneumoniae cells

*Klebsiella pneumoniae* competent cells were generated as previously described (47), with individual colonies containing the pSim6 plasmid inoculated in 5 mL LB media (with 100 µg/mL ampicillin) overnight at 30 °C. Overnight cultures were diluted 1/100 in low salt LB media and incubated at 30 °C to OD_600nm_ = 0.4 - 0.5, with the plasmid activated at 42° C for 1 h. Cells were washed and reconstituted in ice-cold, sterile 10% glycerol.

### Deletion strain generation and confirmation

The lambda-red recombinase system was used to generate the Δ*tolQ* strain (47, 48). Briefly, the chloramphenicol resistance gene was amplified from pkD3 plasmid with *tolQ* complementary overhangs (Table S1), transformed into competent *K. pneumoniae* with pSim6 via electroporation at 1800 V, and incubated at 37 °C for 3 h. Following recovery, cells were plated on chloramphenicol LB agar plates (17 µg/mL chloramphenicol) and incubated overnight at 37 °C. Successful deletions were screened by PCR; using provided primers (Table S1). Whole Genome Sequencing was performed using 2.67 million reads (SeqCenter, Pittsburgh, US) and compared to the reference genome (accession number FO834906.1) (Table S2).

### TolQ-TolR complex structure prediction using AlphaFold

AlphaFold2 predicted structures of *K. pneumoniae* TolQ (A0A0W7ZZF0) and *K. pneumoniae* TolR (A6T6G4) were searched against the Protein DataBank using the DALI server (http://ekhidna2.biocenter.helsinki.fi/dali/) to identify structural homologs with experimentally resolved structures. Retrieved hits were prioritized by degree of experimental characterization and structural similarity to the predicted oligomeric states of *K. pneumoniae* TolQ. The amino acid sequences of both proteins were used as query in AlphaFold3 (https://alphafoldserver.com/) to generate a predicted TolQ–TolR complex, informed by the oligomeric configurations of known homologs (TolQ homolog UniProt ID: V5VAS0, *Acinetobacter baumannii*). Two models were predicted: phosphorylation modifications at residues S98 and S225; and lacking these post-translational modifications. To assess membrane integration, both models were analyzed using the Orientations of Proteins in Membranes (OPM) database (https://opm.phar.umich.edu), enabling prediction of their spatial orientation within the inner membrane of *K. pneumoniae*. Structural visualization and comparative analysis were performed using PyMOL (v3.1.5.1).

### Growth curves

WT and Δ*tolQ* were grown in 100 µL of LB, LM, LM + 10 µM Fe_2_(SO_4_)_3_ and LM + 10 µM ZnSO_4_ media for 24 h in a 96-well clear, sterile polystyrene plate. Bacterial cell densities were determined by plating dilution series and OD_600nm_ measurements were performed to generate a standard curve. The cultures were incubated at 37°C with shaking, measuring OD_600nm_ every 15 min (BioTek HM1 plate reader.) Data were collected in biological triplicate and technical duplicate and plotted using GraphPad Prism v10.

### Macrophage infection

Immortalized BALB/c macrophages were infected with *K. pneumoniae* WT and Δ*tolQ* strains as previously described (49). Macrophages were seeded at 5 x 10^4^ cells per well (24-well plate) in Dulbecco’s modified Eagle’s medium (DMEM) for 48 h at 37 °C with 5% CO_2_. At confluence, DMEM media was removed, cells were washed with PBS, and co-cultured at a multiplicity of infection (MOI) of 100:1 (5 x 10^7^ bacteria cells/mL in DMEM without antibiotics) in triplicate at 37 °C and 5% CO_2_ for 90 min; uninfected macrophages were grown as negative controls.

### Lactate dehydrogenase cytotoxicity assay

Macrophage cytotoxicity was measured by released of lactate dehydrogenase (CytoTox 96® Non-Radioactive Cytotoxicity Assay, Promega, G1780) according to the manufacturer instructions. Maximum death was measured in triplicate by lysing uninfected macrophages. Background absorbance was measured using DMEM + 1.2% triton. Data were plotted and analyzed using GraphPad Prism v10.

### Phagocytosis evasion and enumeration of phagocytosed bacteria

Phagocytic evasion was determined by colony forming unit (CFU) counts of bacteria within the co-culture supernatant. Lysed macrophage debris was plated on LB agar and incubated for 24 h at 37 °C. Phagocytosed bacteria were enumerated by lysing macrophages inoculated with bacteria as described above, serially diluting lysate and plating on LB agar. Plates were incubated for 24 h at 37 °C. Data were plotted and analyzed using GraphPad Prism v10.

### Broth microdilution antibiotic susceptibility assay

Overnight WT and Δ*tolQ* cultures were subcultured at a 1:100 ratio into Cation-adjusted Mueller Hinton Broth (MH). Subcultures were incubated at 37 °C until bacteria reached mid-log phase (approx. 3 h for WT and 5 h for Δ*tolQ*). Cell counts were performed as described above. Biological replicates were diluted to a concentration of 8 x 10^7^ CFUs/mL in MH media with antibiotics: imipenem, ceftriaxone, and ceftazidime at 1 µg/mL – 0.03 µg/mL and meropenem at 250 ng/mL – 1.95 ng/mL. Inoculated 96-well microplates were incubated at 37 °C for 24 h with OD_600nm_ values measured (BioTek HM1 plate reader) to determine minimum inhibitory concentration that prevented at least 90% bacterial growth (MIC_90_).

### Transmission electron microscopy

Overnight cultures of WT and Δ*tolQ* strains were washed with 50 mM HEPES buffer and fixed for 1 h in 2.5% glutaraldehyde and 2% paraformaldehyde. Cells were washed, embedded in noble agar (final concentration, 2%), spread into a thin sheet (1 mm thick), and treated with 1% osmium tetroxide for 45 min. Samples were dehydrated by EtOH, incubated in LR white Resin for 1 h, and transferred and polymerized in LR white overnight at 60 °C. A Reichert-Jung Ultracut E Ultramicrotome was used to prepared 88 nm section, which were collected on formvar-coated grids, stained for 5 min in 5% uranyl acetate and lead citrate for 10 min, and imaged by 120kV field emission transmission electron microscope (FEI Tecnai G2 F20).

### Scanning electron microscopy

Overnight cultures of WT and Δ*tolQ* strains were washed and resuspended in phosphate buffer and adhered onto polished carbon planchets for 30 min at room temperature. Cells were fixed on the planchet using 1.5 mL 2% glutaraldehyde for 30 min, washed and submerged in 1% osmium tetroxide for 30 min. Samples were washed using an EtOH dehydration series, underwent critical point drying (Denton DCP-1), and sputter-coated with gold and palladium (Denton Desk V TSC; 90 s under argon atmosphere) before visualization by scanning electron microscopy (FEI Quanta FEG 250).

### Compound library screening

Using WT and Δ*tolQ* strains, we conducted a high throughput screen of a small-molecule compound library (SPARC Drug Discovery at The Hospital for Sick Children). The library contained 2,500 drug-like compounds exhibiting extensive pharmacophore coverage and chemical diversity. Briefly, overnight cultures were diluted 1:100 in LB media with 50 µL subcultured into each well of 384-well plates containing pre-dispensed compounds (dissolved in DMSO; final concentration of 10 µM). Plates were incubated for 24 h at 37 °C with shaking at 200 rpm and OD_600nm_ values were measured using a microplate reader. The average growth and standard deviations were calculated for sample and control populations. The assay was repeated in triplicate for hit confirmation of compounds that reduced culture growth by more than 2 standard deviations from the negative control mean in the primary screen. Kanamycin (50 µg/mL) wells were included as positive controls and DMSO as negative vehicle controls. Data were archived and analyzed using the CDD Vault database from Collaborative Drug Discovery (Burlingame, CA. www.collaborativedrug.com).

### Compound growth inhibition assay

WT and Δ*tolQ* strains were cultured overnight, diluted 1:100 into MH media with 1-(4-fluorophenyl)-3-(3-hydroxyphenyl)urea (compound 597) added at 0.625 – 20 µM, serially diluted, in sterile polystyrene 96-well plates and incubated for 24 h with shaking at 37 °C. Untreated strains and uninfected MH media served as positive and negative growth controls, respectively. OD_600nm_ was measured at 24 h and growth was normalized (100% was OD_600nm_ with no compound; 0% was OD_600nm_ with media). This assay was performed in biological quadruplicate and technical duplicate.

### Compound cytotoxicity and CFU counting in macrophage

Macrophages were incubated with compound 597 at 10 and 20 µM for quantification of LDH release to assess potential cytotoxicity. CFU counts for a 90 min-co-culture were performed from supernatant and phagocytosed cells in the presence of compound 597 at 1, 10, or 20 µM.

### Statistics and reproducibility

Comparisons between two or three data sets were performed using unpaired two-tailed Student’s *t*-tests and comparisons between four or more data sets were performed using a one-way ANOVA. Significance annotations were applied as follows: *p < 0.0332, **p < 0.0021, ***p < 0.0002 and ****p < 0.0001.

## Results

### Phosphorylation dynamics are driven by iron-availability in K. pneumoniae

Given the central role of phosphorylation in bacterial adaptation to nutrient conditions (50–53), and our prior findings of elevated kinase abundance in response to altered iron and zinc levels within growth media (54, 55), we profiled the phosphoproteome of *K. pneumoniae* to identify modification events regulated by metal availability. To enhance phosphopeptide recovery, we incorporated chloroform–methanol precipitation, peptide desalting prior to enrichment, milder acidic conditions (pH 2.5), and late-stationary phase sampling as these protocol modifications improve recovery, as previously demonstrated in bacterial systems (20, 21, 40, 56–58). This approach resulted in a nine-fold increase in phosphoprotein identification compared to earlier studies conducted in the same media conditions (52, 53), as demonstrated by the identification of all known phosphoproteins in the glycolysis pathway (Fig. S1), a standard benchmark for phosphoproteome completeness (40, 56).

To investigate the phospho-regulatory dynamics of *K. pneumoniae* in response to variations in iron and zinc availability, we identified 640 phosphopeptides: 574 were class I (>75% localization probability) (Fig. 1a). These matched to 358 phosphoproteins, indicating that 7% of the proteome was phosphorylated at the time of collection, consistent with other *K. pneumoniae* phosphoproteome reports in the literature (56, 59). Serine, threonine and histidine were the most common phosphosites (60%, 17% and 13%, respectively), in agreement with other established bacterial phosphoproteome profiles (20, 40, 59). A principal component analysis (PCA) revealed that media conditions accounted for the majority of sample variation (iron-replete replicates clustered separately from zinc-replete and LM; Component 1; 59.93%), with replicate variability contributing 8.67% (Component 2) (Fig. 1b).

**Figure 1:**
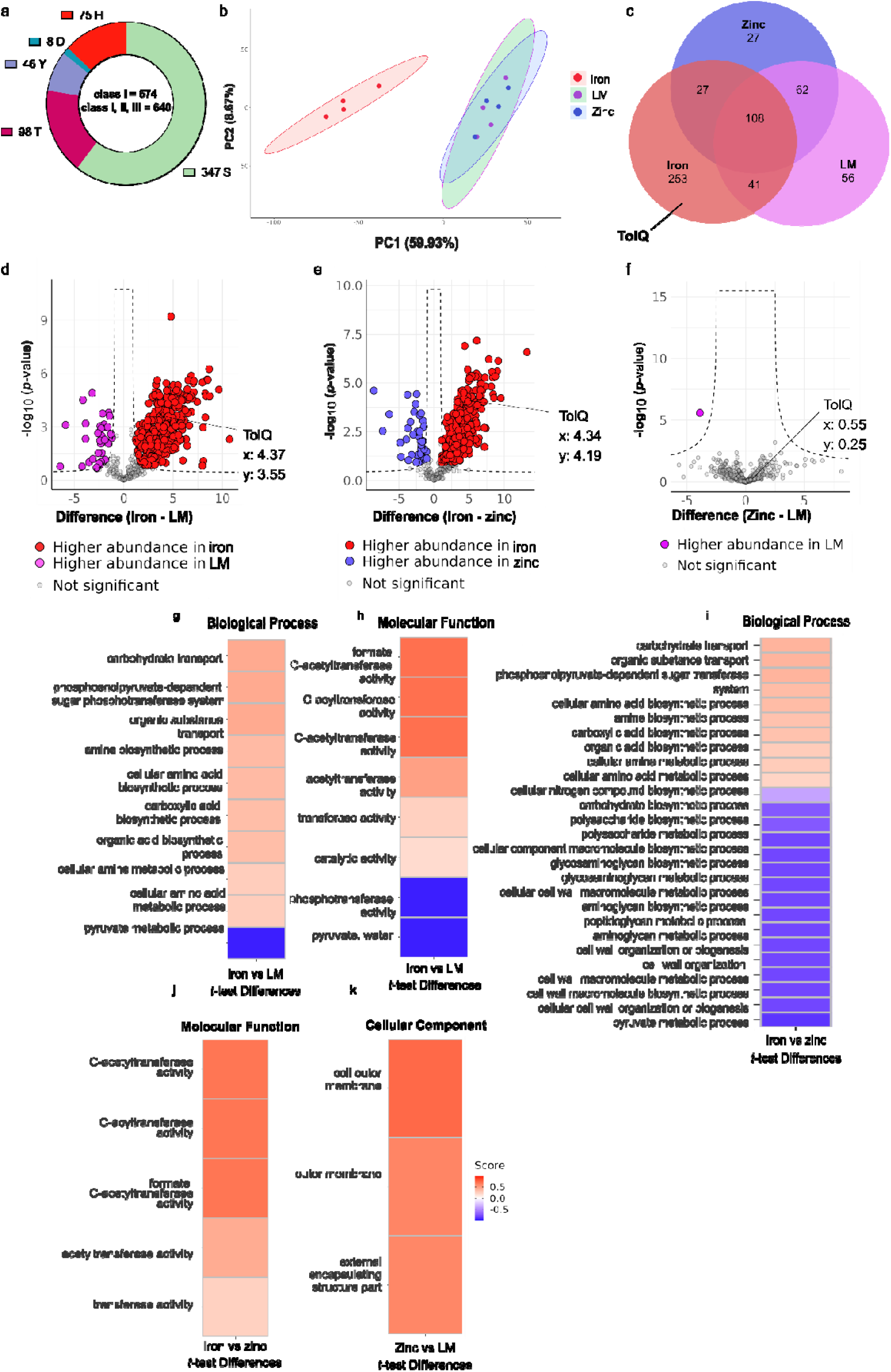
*K. pneumoniae* Phosphoproteome profiling in iron, zinc-replete media and LM. **(a)** phosphosite amino acid distribution identified across all media conditions. **(b)** Principal component analysis (PCA) of sample variability across iron (red), zinc (blue)-replete media and limited media (LM; pink). **(c)** Venn diagram distribution of phosphopeptides identified in iron (red), zinc (blue)-replete media and LM (pink). **(d)** Volcano plot of phosphopeptides significantly more abundant in iron-replete (red) vs LM (purple). **(e)** Volcano plot of phosphopeptides significantly more abundant in iron (red) vs zinc-replete (blue). **(f)** Volcano plot of phosphopeptides significantly more abundant in zinc replete vs LM (purple). **(g)** One-dimensional (1D) UniProt annotation enrichment heatmap of phosphopeptides from GOBP in iron-replete vs LM **(h)** 1D heatmap of GOMF identified in iron-replete vs LM. **(i)** 1D heatmap of GOBP enrichment in iron vs zinc-replete media. **(j)** 1D heatmap of GOMF enrichment in iron vs zinc-replete media. **(k)** 1D heatmap of cellular GOCC in iron vs zinc-replete media. Scores approaching 0.5 (red) indicate that the term is overrepresented in zinc-replete compared to LM, whereas scores approaching −0.5 (blue) indicate that the term is underrepresented. Significance was determined using a Student’s *t*-test with *p*-value ≤ 0.05. FDR = 5% and S_0_ = 1.

To characterize phosphopeptide distribution across media conditions, we generated a Venn diagram to illustrate that iron-replete conditions yielded the highest number of unique phosphoproteins (253), followed by LM (56) and zinc-replete media (27) (Fig. 1c), underscoring iron’s dominant role in regulating phosphorylation-dependent cellular activity. Phosphopeptides that were exclusive to iron-replete conditions were mainly from proteins involved in metabolism (i.e., SerC(60), YqhD(61), YqiE(62), ArgD(63)), cell growth and protein folding (i.e., ClpB(64)), and cell division/membrane stabilization (i.e., TolQ(64), ZapB(65), FtsZ(65)). Increased phosphorylation in iron-replete conditions was further supported by the 10-fold higher abundance of phosphopeptides identified in iron-replete versus LM (362 versus 35, respectively) (Fig. 1d) and nine-fold higher phosphopeptide abundance for iron-replete versus zinc-replete conditions (366 versus 42, respectively) (Fig. 1e). Among the most highly enriched phosphopeptides in iron-replete media were those from biosynthetic enzymes, such as TrpC(66) (83-fold), SerA(67) (725-fold), and GltD(68) (423-fold). TolQ(30), involved in membrane stabilization, exhibited a 20-fold increase in phosphorylation abundance compared to imputed values in iron-limited conditions. Conversely, phosphopeptides enriched under iron-limited conditions compared to iron-replete included those associated with cell shape (e.g., YhcB(69), 2-fold increase), cell wall integrity (e.g., MrcB(70), 11-fold increase), and transport functions (71–77) (e.g., TauC, 14-fold; MglA, 4-fold; PutP, 5-fold; YliA, 5-fold; PrlA, 6-fold; SecD, 7-fold; OmpC, 3-fold). Notably, the abundance of only one protein (RodZ, a cytoskeletal protein involved in cell shape regulation (78)) was significantly different upon comparison of zinc-replete and LM conditions (Fig. 1f). These findings suggest a strong link between iron availability and cellular regulation via phosphorylation, whereby iron initiates the regulation of processes involved in cell growth and the absence of iron may play a role in morphological adaptation, nutrient acquisition, and stress response.

### Phosphoproteome mapping reveals strong connections between iron availability and biosynthesis versus zinc availability and capsule regulation

To identify functional enrichment patterns in our dataset, we performed 1D annotation analysis using UniProt Gene Ontology (GO) terms (79). For GO biological processes (GOBP), we observed that biosynthetic processes, such as amine, carboxylic acid, organic acid and amino acid synthesis, as well as carbohydrate and organic substance transport were more abundant in iron-replete conditions compared to LM (Fig. 1g). For GO molecular function (GOMF), we observed processes, such as acetyltransferase and catalytic activity, that were more abundant in iron-replete conditions compared to LM (Fig. 1h). Conversely, pyruvate metabolic processes and phosphotransferase were enriched in LM conditions. Similarly, the comparison of iron- and zinc-replete conditions, GOBP showed enrichment of biosynthetic pathways, including amino acid, carbohydrate, organic acid, and nitrogen compound biosynthesis in iron-replete conditions compared to zinc-replete (Fig. 1i). For zinc-replete conditions by GOBP, we observed an enrichment in cell wall-associated phosphoproteins, including those involved in polysaccharide, glycosaminoglycan, aminoglycan biosynthesis, and cell wall organization (i.e., *mraY* (80)*, mrcB* (70)*, ompA* (81)*, nagZ* (*82*)*, lpp* (83), and *glmU* (84)). By GOMF, we observed enrichment of acetyltransferase activity in iron-replete conditions with no enrichment under zinc-replete conditions (Fig. 1j). Furthermore, 1D annotation of subcellular localization by GO cellular component (GOCC) revealed enrichment of outer membrane and capsule proteins in zinc-replete media, reinforcing the notion of enhanced cell envelope regulation under zinc stress (Fig. 1k). Collectively, these findings highlight iron-replete conditions promoting growth-associated phosphorylation events, whereas zinc-replete and LM conditions elicit stress-associated phosphorylation patterns, particularly in energy metabolism and cell envelope regulation.

### Computational modeling predicts that phosphorylation causes major conformation changes in the TolQ-TolR complex, influencing iron homeostasis and defining a putative broad-spectrum antimicrobial target

Given the exclusive phosphorylation of TolQ in iron-replete conditions and its established role in cell growth and membrane stabilization (31), we performed computational modeling to predict the structural impact of phosphorylation, identify interacting partners, and assess evolutionary conservation. Through this approach, we aimed to elucidate the connection between iron availability, phosphoregulation, and cell growth, and evaluate TolQ’s potential as a broad-spectrum antimicrobial target in Gram-negative bacteria. First, we confirmed phosphorylation of TolQ in iron-replete conditions via manual inspection of MS/MS fragmentation spectra, which revealed phosphorylation at two serine residues. Phosphorylation of S98 was confirmed by characteristic H□PO□neutral losses (97.9769 amu) observed in the doubly charged y_13_ and y_14_ ions, and in singly charged b_3_, b_4_, b_5_, and b_7_ (Fig. 2a). Notably, the b_9_ fragment exhibited a mass shift of +79.96 amu relative to its theoretical unmodified *m/z*, indicating the presence of an HPO_4_ group within the fragment. Similarly, S225 phosphorylation was validated by H PO neutral losses observed in y_4_ – y_9_ ions (Fig. 2b).

**Figure 2:**
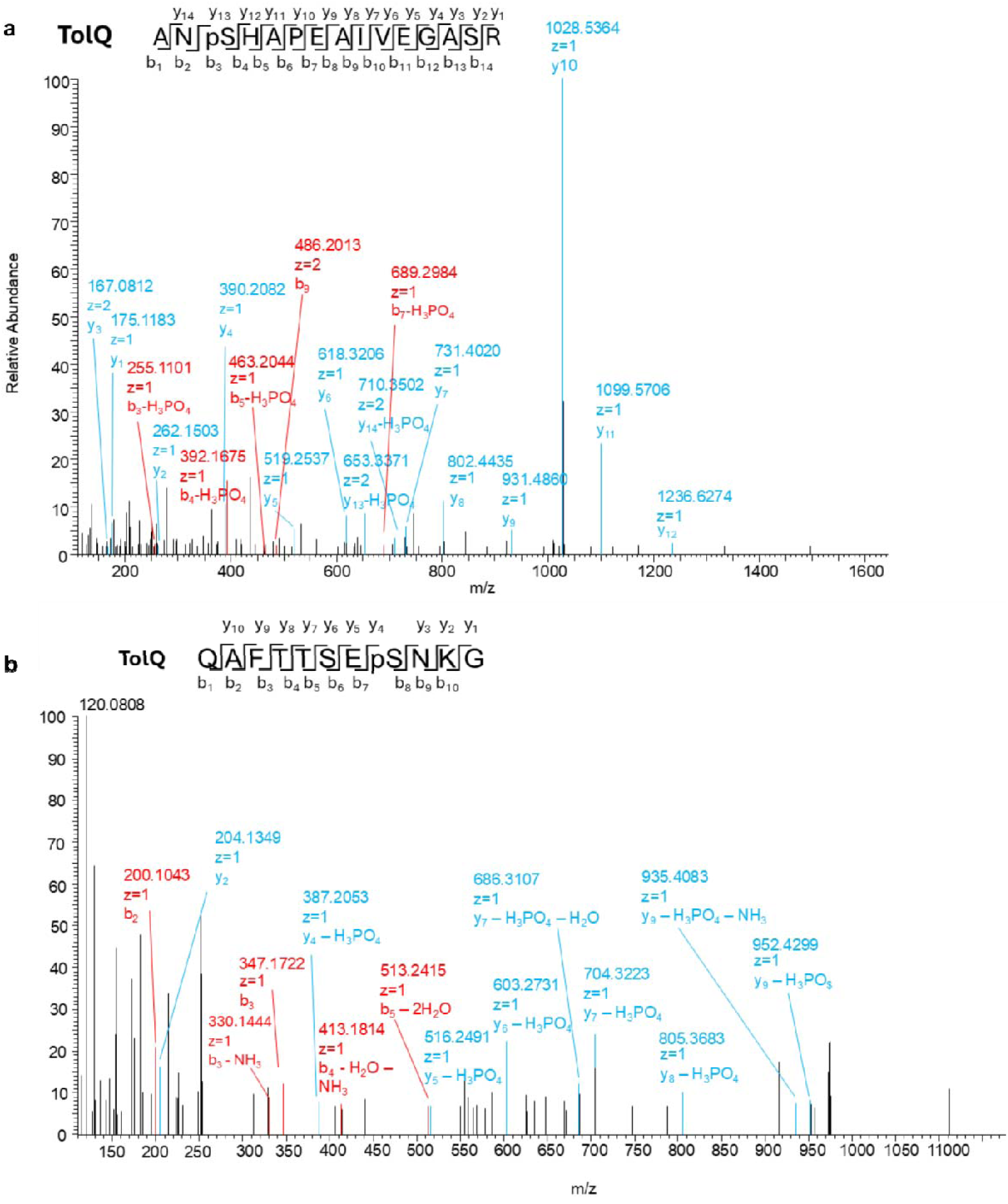
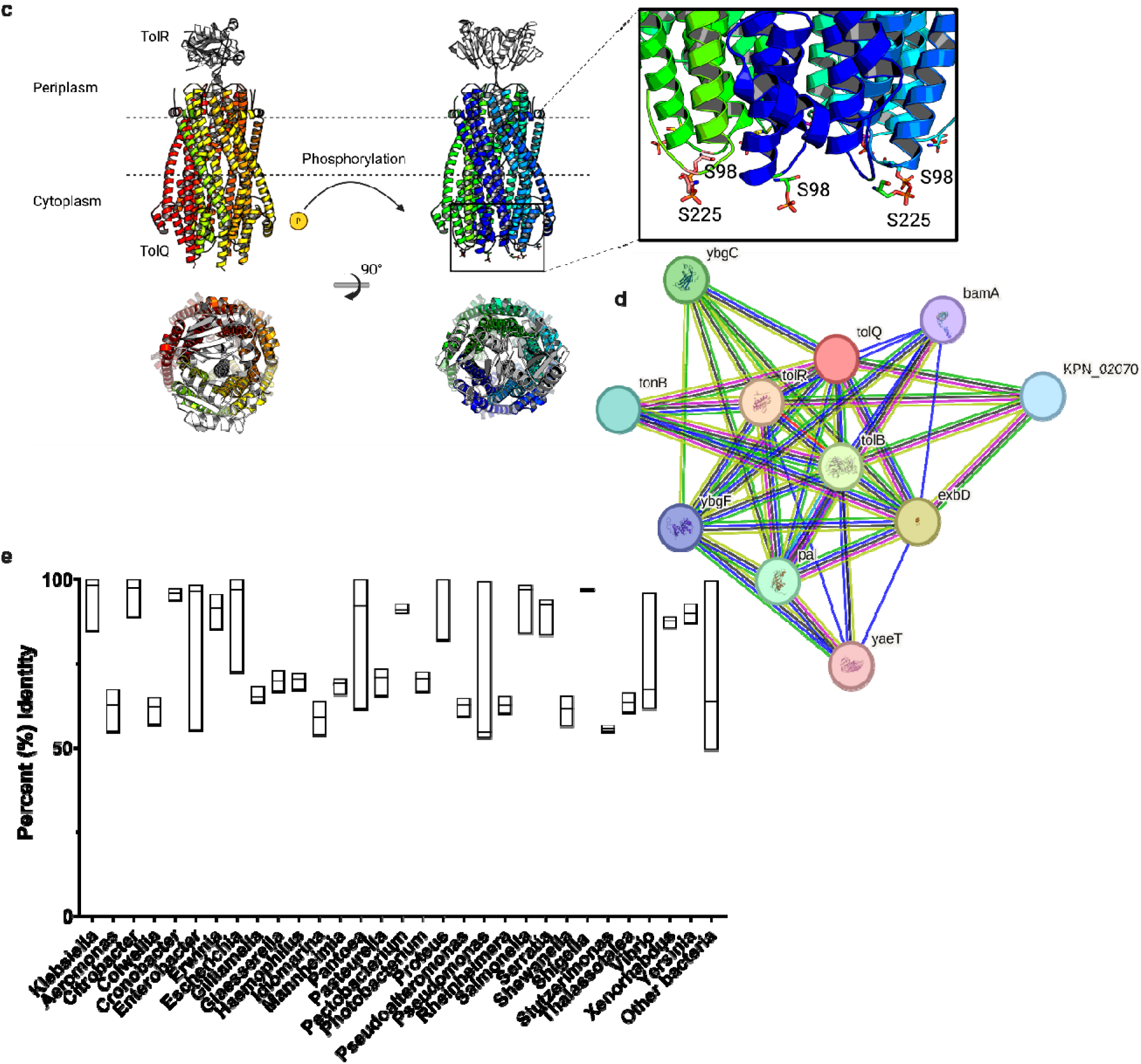
Bioinformatic characterization of *K. pneumoniae* TolQ protein. **(a)** MS/MS spectra of ANpSHAPEAIVEGASR phosphopeptide corresponding to S98 phosphosite identified in iron-replete condition. **(b)** MS/MS spectra of QAFTTSEpSNKG phosphopeptide corresponding to S225 phosphosite identified in iron-replete condition. **(c)** AlphaFold2 structure prediction with highlighted S98 and S225 phosphosites. Designed in PyMOL. **(d)** Protein-protein interaction network from STRING database. **(e)** Percent of shared sequence identity between bacterial species for TolQ based on BLASTp search. Maximum number of target species was set to 5,000.

Next, to investigate the predicted structural impact of phosphorylation of TolQ, we conducted a Protein Data Bank (PDB) (85) search via the DALI (Distance-matrix ALIgnment) (86) server to identify structurally homologous proteins with validated crystal structures. Candidate homologs were prioritized based on their degree of experimental characterization and root mean squared deviation (RMSD) values across oligomeric states. *A. baumannii* TolQ was selected due to its well-characterized structure (30), 47% sequence identity with *K. pneumoniae* TolQ, and a high overlap of predicted and resolved structure (RMSD of 1.054 Å; Fig. S2). This RMSD value (< 2 Å) indicates a high-confidence structural alignment between the AlphaFold-predicted model and the experimentally resolved *A. baumannii* TolQ structure (87). Searching of the OPM database(88) revealed that both phosphorylated serines were intracellular-facing and AlphaFold (89) predictions indicated that phosphorylation at Ser98 and Ser225 induced conformational changes in the TolQ-TolR complex, via a 90° rotation of TolR relative to TolQ (Fig. 2c). The predicted template modeling scores (pTM) were 0.77 for the phosphorylated complex and 0.75 for the unphosphorylated form, indicating high structural confidence for both states (90, 91).

To better understand the role of iron in TolQ regulation, we used the STRING (92) database to identify potential TolQ interaction partners. We observed predicted interactions with core components of the Tol-Pal system (TolA, TolB, TolR, Pal), as well as proteins involved in membrane integrity and assembly: *ybgC* (phospholipid metabolism) (93), *ybgF* (peptidoglycan synthesis and outer membrane constriction) (94), and *bamA* and *yaeT* (outer membrane protein assembly) (95, 96) (Fig. 2d). Notably, TolQ also interacts TonB and ExbD, key components of the Ton system, which couples free energy from the proton motive force with iron transport across the outer membrane (97, 98), providing a connection with iron homeostasis. Further, to evaluate the conservation of TolQ and its potential as a broad-spectrum antimicrobial target, we performed a BLASTp homology search using the full-length TolQ amino acid sequence against 5,000 bacterial species spanning over 30 genera. The top 5,000 hits were selected based on bit score (reflecting alignment quality via evolutionarily plausible substitutions and alignment gap penalties), E-value (indicating the statistical likelihood of the alignment occurring by chance), percent identity (amino acid sequence similarity), and query coverage (extent of the query sequence aligned) (99, 100). TolQ exhibited >50% sequence identity in 99.96% of species queried, with 1,612 species showing >90% identity (Fig. 2e). All sequences exceeded the 30% identity threshold commonly used to infer homology and potential structural, conformational, and functional similarity (100–102) while many surpassed the 90% threshold indicative of near-identical proteins with minor mutations(103). Notably, many genera with conserved TolQ homologs include clinically relevant pathogens associated with antibiotic resistance and human morbidity or mortality, such as *Salmonella*, *Shigella*, *Escherichia*, *Pseudomonas*, *Enterobacter*, *Citrobacter*, *Cronobacter*, *Proteus*, *Haemophilus*, *Yersinia*, *Vibrio*, and *Serratia* (104–107). Together, these findings suggest that phosphorylation modulates TolQ-TolR conformation, with implications for membrane stability and cell division, and illustrates an interaction network between TolQ and the iron acquisition TonB system. The widespread conservation and functional centrality of TolQ in a membrane stabilization network suggests that it may be strategically targeted to disrupt bacterial membranes across diverse Gram-negative pathogens, offering a promising avenue for antimicrobial development.

### ***Δ***tolQ strains display reduced growth and viability with increased susceptibility to ***β***-lactams and host immune clearance

Given TolQ’s role in the membrane stabilization network of *K. pneumoniae*, we hypothesized that its disruption could impair bacterial growth, virulence, and resistance to β-lactam antibiotics targeting the cell envelope. To test this, we conducted growth curve analyses in LB and iron-supplemented media (i.e., conditions under which TolQ is phosphorylated) alongside minimum inhibitory concentration (MIC) broth microdilution assays using β-lactams, cytotoxicity assays, and CFU counts to assess virulence. Growth curve data revealed that Δ*tolQ* entered the exponential phase 2-3 hours later than WT strains in both LB and iron-supplemented media (Fig. 3a, c). The area under the curve (AUC) for WT was 27% greater than Δ*tolQ* in LB (p < 0.0001; Fig. 3b) and 21% greater in iron-supplemented media (p < 0.0001; Fig. 3d). Next, given TolQ’s role in stabilizing the periplasmic space (30) (i.e., location of the peptidoglycan layer targeted by β-lactams (13, 14)), we predicted that Δ*tolQ* would exhibit increased β-lactam susceptibility. To test this hypothesis, a broth microdilution MIC assay was performed in MH media to identify the minimum concentration of antibiotic which reduced WT and Δ*tolQ* growth by ≥ 90%. Δ*tolQ* strains showed a two-fold increase in susceptibility to imipenem, meropenem, and ceftazidime, and an eight-fold increase to ceftriaxone (Fig. 3e-g). Further, to investigate whether membrane instability affected virulence and immune clearance, we performed LDH cytotoxicity and CFU assays using co-cultures of WT and Δ*tolQ* strains with BALB/c macrophages. LDH assays showed comparable macrophage cell death: WT induced 25% death, while Δ*tolQ* induced 24% (Fig. 3h). Uninfected macrophages exhibited 15% death, indicating ∼10% increased death due to bacterial exposure. The increase was statistically significant for WT (p = 0.0063), nearing significant for Δ*tolQ* (p = 0.0503), and not significantly different between the two strains (p = 0.8683). LDH assay results indicated that WT and Δ*tolQ* have similar cytotoxicity profiles; however, Δ*tolQ* strains were significantly more susceptible to phagocytosis. After 90 minutes of co-culture, lysed infected macrophages showed 5.5-fold higher Δ*tolQ* CFU (mean = 7,593) than WT (mean = 1,371; p = 0.0280; Fig. 3i), indicating increased phagocytosis of Δ*tolQ* compared to WT. Additionally, the supernatant from Δ*tolQ* co-cultures yielded three-fold fewer colonies than WT (p = 0.0003; Fig. 3j), confirming increased phagocytosis of Δ*tolQ* by macrophages. Together, phenotypic characterization of Δ*tolQ* underscores its potential as an antimicrobial target. Δ*tolQ* exhibited impaired growth, heightened β-lactam susceptibility, and diminished evasion of phagocytosis. These findings collectively demonstrate that TolQ disruption attenuates *K. pneumoniae* and may enhance the therapeutic efficacy of existing antibiotics.

**Figure 3:**
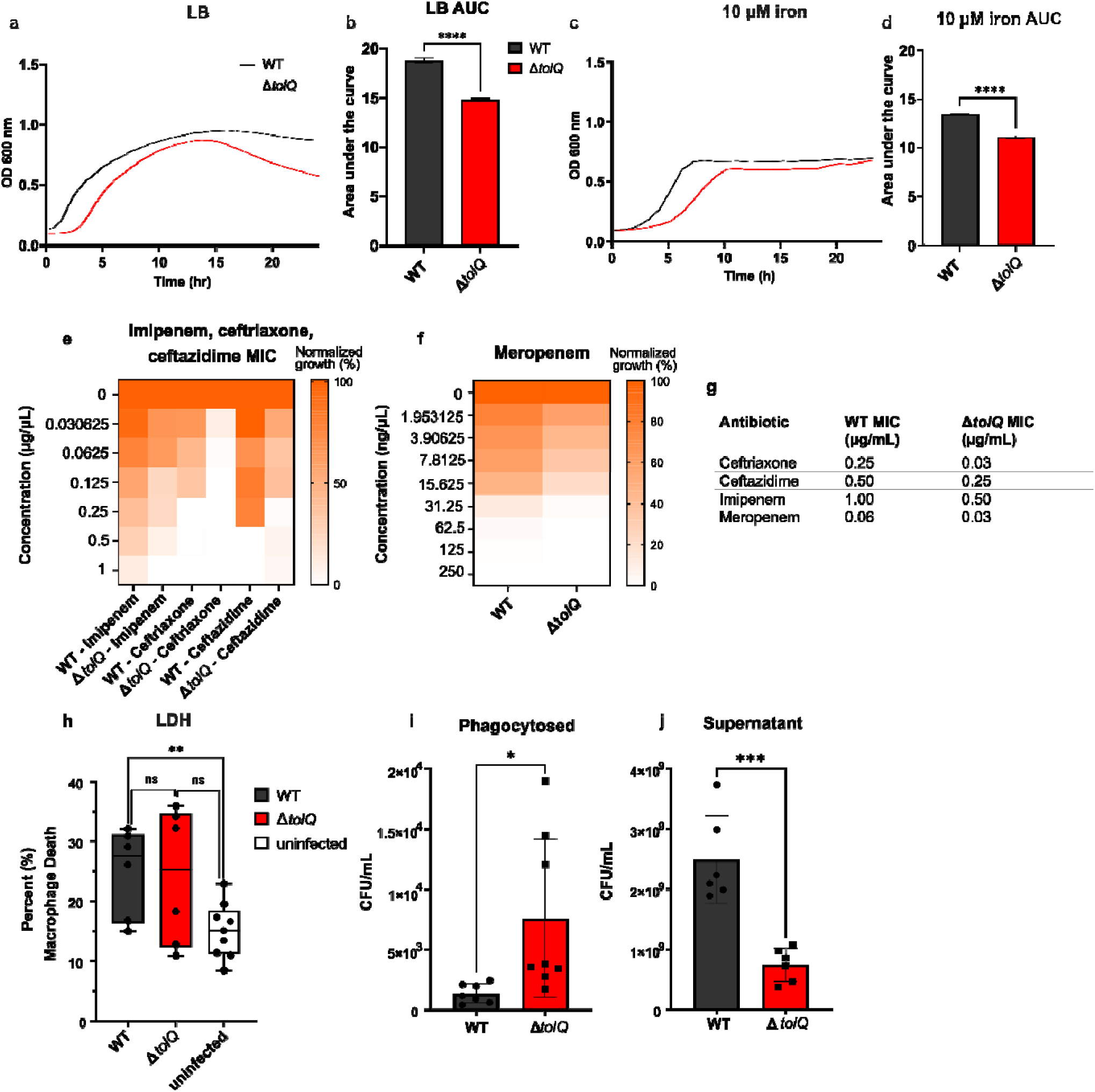
Phenotypic characterization for WT (black) and Δ*tolQ* (red) *K. pneumoniae*. **(a)** Growth curve in nutrient rich (LB) media. **(b)** AUC for nutrient rich LB media growth curve Growth curve in LB media. **(c)** Growth curve in iron supplemented M9 media. **(d)** Area under the curve (AUC) for iron supplemented M9 media. OD_600nm_ measured every 15 min in biological triplicate and technical duplicate. Error bars denote standard error. **(e)** Heatmap depicting minimum inhibitory concentration (MIC) for WT and Δ*tolQ* susceptibility to the β-lactams imipenem, ceftriaxone and ceftazidime**. (f)** Heatmap depicting minimum inhibitory concentration (MIC) for WT and Δ*tolQ* susceptibility to the β-lactam meropenem. **(g)** MIC values for WT and Δ*tolQ* **(h)** Lactate Dehydrogenase (LDH) cytotoxicity assay for macrophages infected with WT or Δ*tolQ K. pneumoniae* vs uninfected macrophage cell death. **(i)** Colony forming units (CFU) for phagocytosed WT and Δ*tolQ K. pneumoniae*. **(j)** CFU for WT and Δ*tolQ* in the supernatant following macrophage co-culture. Statistical difference between strains was calculated using a Two-tailed Student’s t-test where *p < 0.0332, **p < 0.0021, ***p < 0.0002 and ****p < 0.0001.

### Electron microscopy illustrates cell division and membrane disruption in ***Δ***tolQ

To investigate the reduction in cell density observed beyond 15 h in LB media we used transmission electron microscopy (TEM) and scanning electron microscopy (SEM) on overnight cultures of WT and Δ*tolQ* grown in LB media. These images showed that overnight WT cultures were a mix of rod-shaped (Fig. 4a,e) and spherical cells (Fig. 4b), with the majority demonstrating a rod-shaped morphology. Conversely, Δ*tolQ* cells, were exclusively spherical and displayed a “ruffled” membrane (Fig. 4c) compared to the healthier “smooth” membrane of WT. Most images of Δ*tolQ* cultures revealed cells devoid of intracellular contents, consistent with increased lysis due to impaired cell envelope integrity (Fig. 4d,f). This observation is similar to what has been reported for *K. pneumoniae* lysed by ampicillin (108). Δ*tolQ* SEM micrographs also showed a filamentous morphology indicative of impaired cell division, which is consistent with Tol mutants in comparable bacterial species (33) (Fig. 4f). Together, these data highlight differences in cell morphology and viability between WT and Δ*tolQ* strains, with Δ*tolQ* displaying significant membrane instability, aligning with its known functional roles.

**Figure 4:**
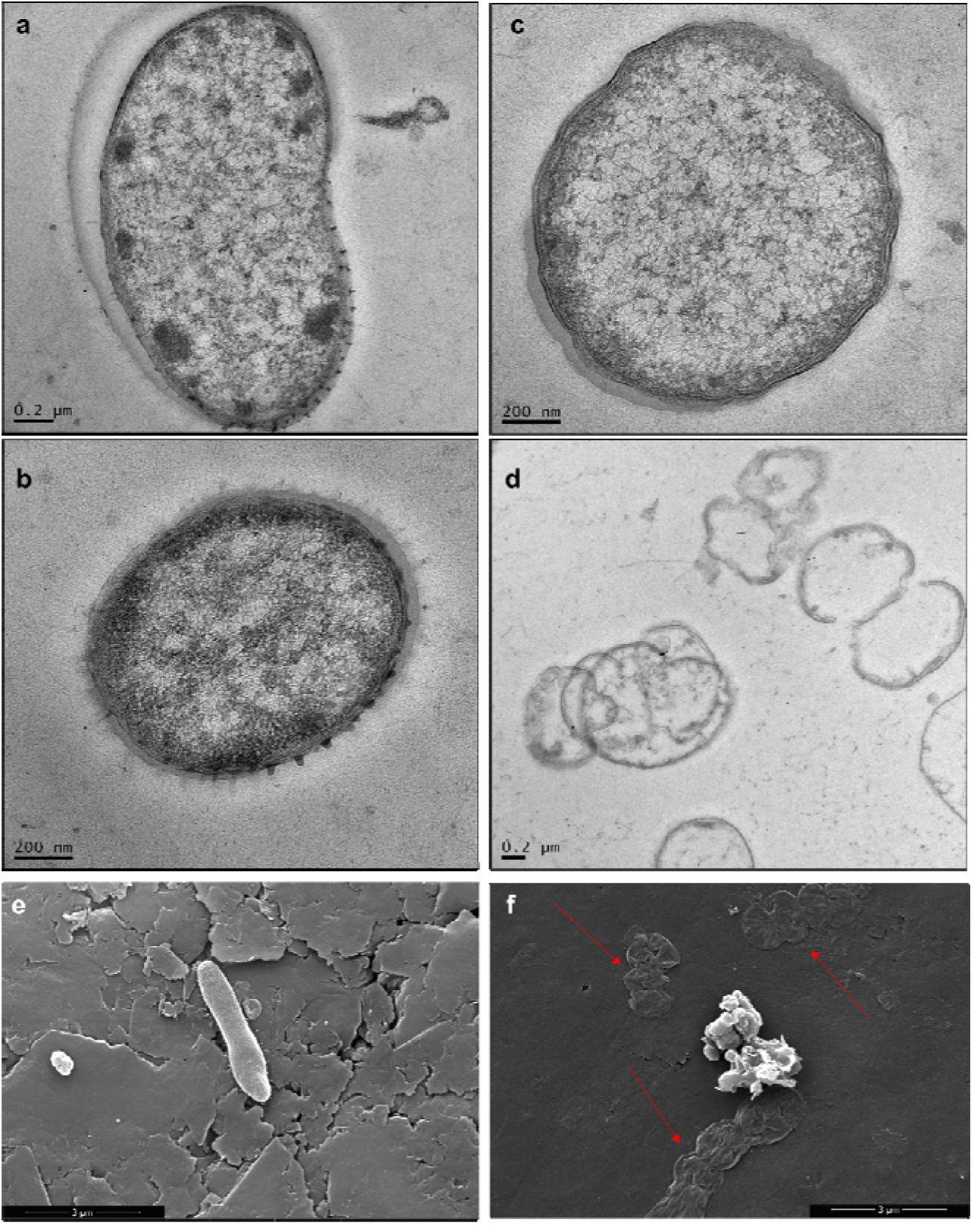
Cell morphology visualization via electron microscopy. **(a)** Transmission electron microscopy (TEM) micrograph of rod-shaped WT cell. **(b)** TEM micrograph of spherical WT cell. **(c)** TEM micrograph of spherical Δ*tolQ* displaying ruffled membrane **(d)** TEM micrograph of devoid Δ*tolQ* cells. Strains were cultured in LB media and collected at stationary phase. **(e)** Scanning electron microscopy (SEM) micrograph of 30,000X magnification of WT cells. **(f)** SEM micrograph of 30,000X magnification Δ*tolQ* illustrating devoid cells. Images are representative of the general morphology from ≥20 captured images from ≥5 fields of view.

### High-throughput drug screening identifies compounds that target TolQ to impair bacterial growth and survival

Following validation of TolQ as a putative antimicrobial target, we sought to identify compounds that selectively inhibit WT but not Δ*tolQ*, which would indicate TolQ-specific activity. Both strains were incubated with a compound library of 2,500 drug-like compounds. Compounds that reduced OD_600nm_ measurements by at least two standard deviations compared to untreated samples were characterized as ‘hits’ (Fig. S3). From this screen and in combination with CDD Vault’s pattern recognition algorithms, we prioritized 1-(4-fluorophenyl)-3-(3-hydroxyphenyl)urea (compound 597; Fig. 5a). To assess the effects of compound 597 on *K. pneumoniae* growth, we performed broth microdilution assays in MH media. We observed significant inhibition of WT growth at 20 µM (Fig. 5b), while growth of the Δ*tolQ* strains remained unaffected (Fig. 5c).

**Figure 5:**
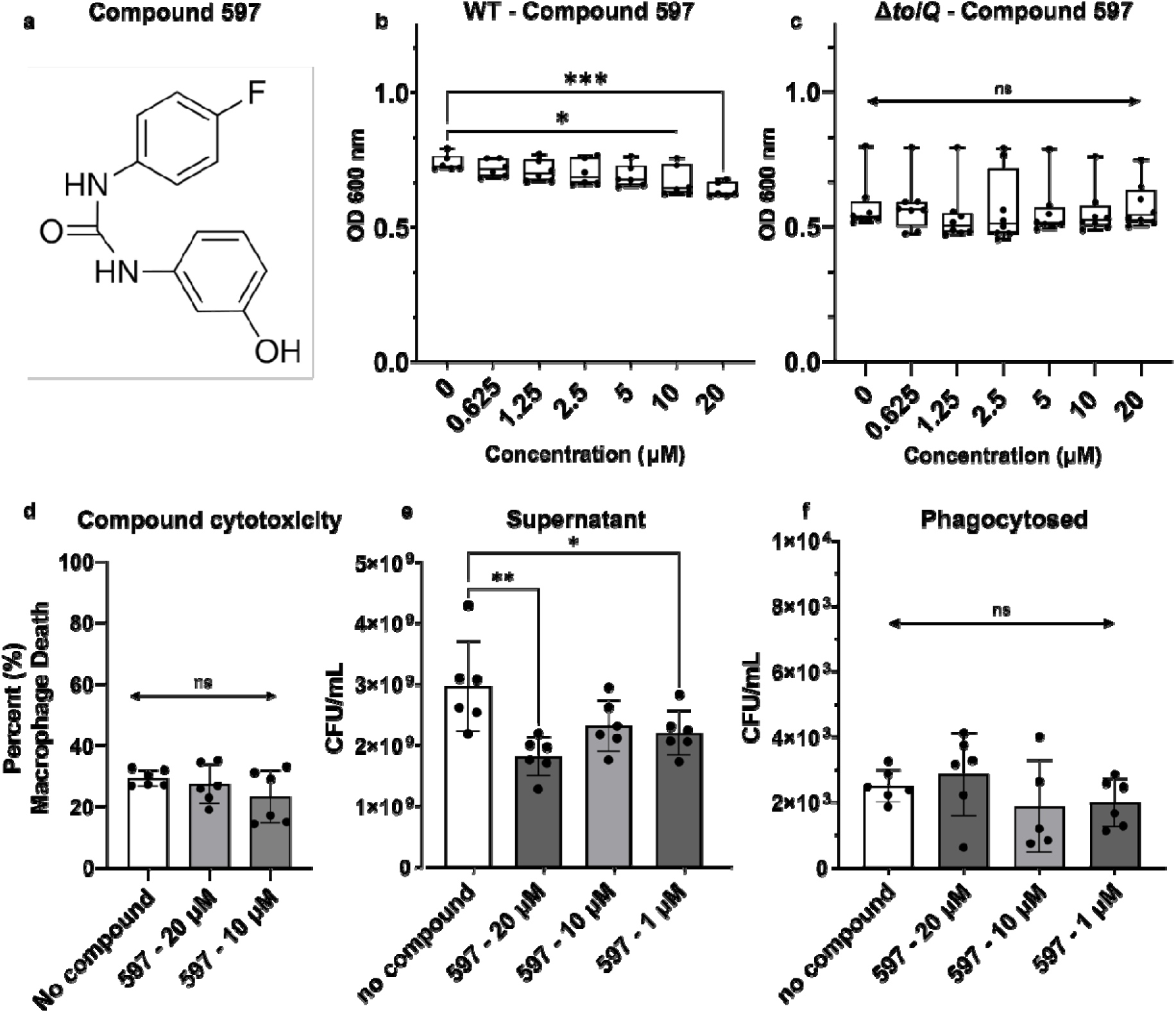
High-throughput drug screening identification and characterization of compound that affects WT but not Δ*tolQ K. pneumoniae*. (**a**) Molecular structure of biosimilar 1-(4-fluorophenyl)-3-(3-hydroxyphenyl)urea (compound 597). (**b**) OD_600nm_ of WT in increasing concentration of compound 597. (**c**) OD_600nm_ of Δ*tolQ* in increasing concentration of compound 597. Box and whisker plot show the median, second and third quartile and minimum and maximum values. (**d**) LDH assay showing percent macrophage death in the presence of compound 597 compared to untreated. (**e**) WT supernatant CFU following 90 min co-culture with BALB/c macrophages in the presence of increasing concentrations of compound 597. (**f**) BALB/c macrophage phagocytosed WT CFU in the presence of increasing concentrations of compound 597 during 90 min infection. Error bars denote standard deviation. Statistics were performed using One-way ANOVA, where *p < 0.0332, **p < 0.0021, and ***p < 0.0002.

Next, we hypothesized that WT *K. pneumoniae* treated with compound 597 would phenocopy Δ*tolQ* upon co-culture with macrophages. We first performed an LDH assay on macrophages exposed to increasing concentrations of compound 597 with no significant cytotoxicity observed at the tested concentrations (p-values at 20 µM = 0.9489, 10 µM = 0.2860) (Fig. 5d). We also conducted infection assays using WT *K. pneumoniae* co-cultured with BALB/c macrophages in the presence or absence of compound 597 and observed a statistically significant reduction in CFU recovered from the supernatant when the compound was present, suggesting enhanced bacterial clearance (Fig. 5e). Specifically, compound 597 showed reduced bacterial recovery from the supernatant of 39% at 20 µM (p = 0.0003), 22% at 10 µM (p = 0.0675), and 26% at 1 µM (p = 0.0227). These reductions mirror the enhanced phagocytosis observed in Δ*tolQ*, although the magnitude of effect was less pronounced than the three-fold decrease in supernatant CFU seen with Δ*tolQ*. Since these compounds only moderately decrease bacterial growth (9-17%), the greater reduction in CFU likely reflects increased phagocytosis. Interestingly, CFU counts from lysed macrophages did not differ significantly between untreated WT and compound-treated conditions (Fig. 5f); however, visually there does appear to be an increase in CFU for compound 597 at 20 µM compared to untreated *K. pneumoniae* (14% increase, p = 0.9948). Notably, we observed a similar pattern in supernatant reduction and maintained CFU counts from lysed macrophage for Δ*sbmA*(35). These data may suggest that compound-treated bacteria are more effectively phagocytosed and cleared within the phagolysosome, resulting in fewer viable colonies post-co-culture.

## Discussion

This study provides phosphoproteome profiling of *K. pneumoniae* under limited metal ion availability to explore the connection between iron and zinc phosphoregulation and bacterial susceptibility to antimicrobials. To enhance phosphopeptide recovery, we incorporated several method refinements, which increased phosphoprotein identification by nine-fold compared to previously published results using an unmodified method (36, 37). Enhanced phosphopeptide enrichment yielded broader phosphoproteome coverage, expanding the pool of candidates involved in key cellular processes and enabling their selection for further characterization as potential targets for disruption. Specifically, given exclusive phosphorylation in iron-replete conditions and an established role in cell growth and membrane stabilization (31), along with a coordinated role in the Tol-Pal system, we prioritized TolQ as a candidate target for disruption and therapeutic intervention within the present study. Our characterization of TolQ explored a connection with phagocytic evasion, which is supported by our previous study that uncovered a role for activation or iron acquisition systems in phagocytic evasion in *K. pneumoniae* and macrophage co-culture, which correlates with the observed iron-response phosphoregulation of TolQ with modulatory roles in bacterial survivability (109).

Our computational analysis revealed that the phosphopeptides identified in LM and zinc-replete growth conditions clustered together, whereas those identified in the iron-replete condition clustered separately, indicating that phosphorylation dynamics were most influenced by the presence versus absence of iron. Iron-replete conditions resulted in the most phosphorylation events overall, indicating enhanced cellular activity occurring in iron-replete conditions as evidenced by increased regulatory events. The phosphopeptides identified in the iron-replete conditions were from proteins involved in a diverse array of metabolic and biosynthetic processes, further illustrating the importance of this metal nutrient for the bacteria. Of particular interest was the phosphorylation of acetyltransferases in iron-replete conditions since there has not been an established link between iron availability and acetyltransferase phosphor-regulation in *K. pneumoniae*. This illustrates the dynamic and complex interactions of regulatory processes within bacteria, as phosphorylation appears to regulate the activity of acetyltransferases, which regulate the activity of proteins through acetyl groups. Protein acetylation often occurs in metabolic processes in bacteria (i.e., acetylation of coenzyme A) (110). This correlates strongly with 1D annotation results we saw in iron-replete conditions, where there was an enrichment of amine metabolic activity. The effect of iron on the “acetyl-proteome” of *K. pneumoniae* in response to iron warrants exploration due to the parallel importance of iron and acetyltransferases in host-pathogen interaction (111).

Minimal media and zinc-replete media conditions resulted in the phosphorylation of proteins associated with energy generation and cell envelope formation. This may reflect a stress response, as cells attempt to repair or reinforce the envelope in the face of zinc-induced membrane damage. These findings align with previous observations that zinc, released from ZnO nanoparticles, disrupts bacterial cell membrane via electrostatic forces (112). Phosphoregulation of envelope proteins in response to zinc may therefore, represent an adaptive mechanism to counteract metal induced stress. Targeting these proteins could compromise bacterial survival under envelope stress or disrupt envelope organization entirely. For example, Penicillin□binding protein (PBP) 1B (*mrcB*) was among the most abundantly phosphorylated proteins under zinc□replete conditions; this enzyme is a well□established target of β□lactam antibiotics, one of the most widely prescribed antibiotic classes (113). These findings highlight the potential of phosphoproteomic profiling to uncover clinically relevant vulnerabilities and suggest that other zinc□regulated proteins identified here warrant further investigation for their roles in envelope stabilization and antimicrobial susceptibility.

We identified that the inner membrane protein, TolQ, was phosphorylated exclusively under iron-replete media conditions, establishing a link between iron availability and TolQ regulation. We hypothesized that this phosphorylation connects to the role of the Tol-Pal system in membrane stabilization during cell division, a process that is activated when iron is abundant. TolQ phosphosites (S98 and S225) were predicted to reside within intracellular domains, which are commonly phosphorylated to mediate signal transduction and modulate protein conformation and activity (114). Structural predictions using AlphaFold suggested that phosphorylation at these sites induces a 90° rotation of its interacting partner, TolR. The rotation of TolR following TolQ phosphorylation could serve several potential purposes, including the generation of torque, analogous to the mechanisms observed in bacterial flagellar systems (115). Consistent with this, TolQ shares homology with MotA (116), a critical component of the flagellar stator that generates torque, supporting the idea that TolQ phosphorylation may drive torque through TolQ-TolR interactions. Rotary force has previously been proposed to facilitate TolQ-TolR-TolA engagement with the TolB-Pal complex (117). In addition, TolR rotation could influence its interaction with the peptidoglycan layer. Molecular modeling studies have shown that rotation of TolR relative to TolQ produces “open” and “closed” conformations, with only the open state capable of binding peptidoglycan via electrostatic interactions (118). Phosphorylation may therefore facilitate transitions between these conformations, modulating TolR’s interaction with the cell wall. Further experimental evidence will be required to validate the precise role of phosphorylation in TolR function.

Conservation analysis underscored the central role of TolQ in membrane stabilization and revealed high conservation and ubiquity across Gram-negative species, identifying it as a promising broad-spectrum antimicrobial target. To explore this observation, we characterized Δ*tolQ* and observed reduced growth, spontaneous cell lysis, and diminished virulence, supporting TolQ as a potential therapeutic target. We hypothesized that Δ*tolQ* strains were particularly vulnerable to β-lactam antibiotics, including meropenem, imipenem, ceftazidime, and ceftriaxone, due to its function in maintaining cell envelope integrity. Disruption of TolQ likely compromises outer membrane stability, enhancing antibiotic uptake. Given that β-lactams target peptidoglycan synthesis within the periplasm (119), TolQ disruption may exacerbate cell envelope vulnerabilities, amplifying the bactericidal effects of these agents. Interestingly, PBP 1B colocalizes with Tol proteins at the division septum (94), suggesting that perturbation of either component may impair cell division. Notably, Δ*tolQ* exhibited an 8-fold increase in susceptibility to ceftriaxone, compared to a 2-fold increase for the other β-lactams tested. This heightened sensitivity is unexpected, given ceftriaxone’s shared mechanism of action with other β-lactams, namely, inhibition of PBP3-mediated cell wall synthesis (13, 14). It is possible that ceftriaxone more effectively exploits membrane instability in Δ*tolQ* strains or crosses the compromised outer membrane more efficiently. To explore this further, proteomic or transcriptomic comparisons between WT and Δ*tolQ* strains could reveal differential abundance of porins, efflux pumps, or resistance genes that may explain ceftriaxone’s enhanced activity. These findings demonstrate that targeting TolQ potentiates β-lactam efficacy and offers a strategy to extend the clinical utility of these antibiotics, a critical need amid rising β-lactam resistance in *K. pneumoniae* (120).

## Conclusion

This study defines iron-responsive phosphorylation as a key regulatory mechanism underlying bacterial adaptation to nutrient availability in *Klebsiella pneumoniae*. Through integrated phosphoproteomics, functional characterization, and chemical screening, we identify TolQ with regulatory roles of cell envelope integrity and bacterial fitness for targeted disruption. Our findings reveal a link between iron availability, predicted phosphorylation-dependent modulation of the Tol–Pal system, and susceptibility to host defenses and antibiotic treatment. By uncovering this regulatory vulnerability, this work highlights the potential of targeting phosphoregulated membrane systems to disrupt essential bacterial processes and advance the development of new antimicrobial strategies against drug-resistant pathogens.

## Author Contributions

C.R. & J.G.-M. conceptualized the study. C.R., J.S., & J.G.-M. designed the study. C.R., J.S., S.C & N.C. performed experiments and data analysis. C.R. & O.R. designed and developed figures. C.R. & J.G.-M. wrote and edited the manuscript. All authors have read and approved the submitted manuscript.

## Acknowledgments

Thank you to Rapid Novor for allowing the use of their mass spectrometers, SPARC Drug Discovery at The Hospital for Sick Children for assistance with high-throughput compound screening and the Molecular and Cellular Imaging Facility and the University of Guelph for assisting with electron microscopy sample preparation and imaging.

## Data Availability

Data are available via ProteomeXchange with identifier PXD072239.

## Supplementary files

**Table S1:** Primer design.

**Table S2:** Whole genome sequencing of Δ*tolQ* compared to WT (FO834906.1 accession).

**Figure S1:** Phosphoproteome profiling with identification of known phosphoproteins in glycolysis. Pathway mapping performed using KEGG pathways.

**Figure S2:** *K. pneumoniae* TolQ (A0A0W7ZZF0) superimposed with *A. baumannii* TolQ.

**Figure S3:** Optical density (OD_600nm_) of *K. pneumoniae* strains following exposure to the drug-like curated compound library.

