## Supplementary material for "Iron-responsive phosphorylation of TolQ modulates cell envelope integrity and antibiotic susceptibility in *Klebsiella pneumoniae*": Supp files

**Table S1: Primer design.**

| Gene of interest | Forward (F)/ Reverse (R) | Primer Sequence |
| --- | --- | --- |
| *tolQ* | Cm-F | TTAATGAAGCCACGTGCGCTTCCCAAGTCTATT  GTCGCGGAGTTTAAGCAGTGTAGGCTGGAGCTGCTTC |
| *tolQ* | Cm-R | ATCTCGGACTTC  AGTTCGCGACGTCCGCGTCCACGTGCTCTG  GCCATGGCCATATGAATATCCTCCTTAG |
| *tolQ* | F-size | GATGCGCGGCACCTCTTTGG |
| *tolQ* | R-size | GTCCGGCAGATCGACTTCCA |
| *tolQ* | R-in/out | GATGCGCGGCACCTCTTTGG |
| *Cat* | F-in/out | GCAACTGACTGAAATGCCTC |

**Table S2: Whole genome sequencing of Δ*tolQ* compared to WT (FO834906.1 accession).**

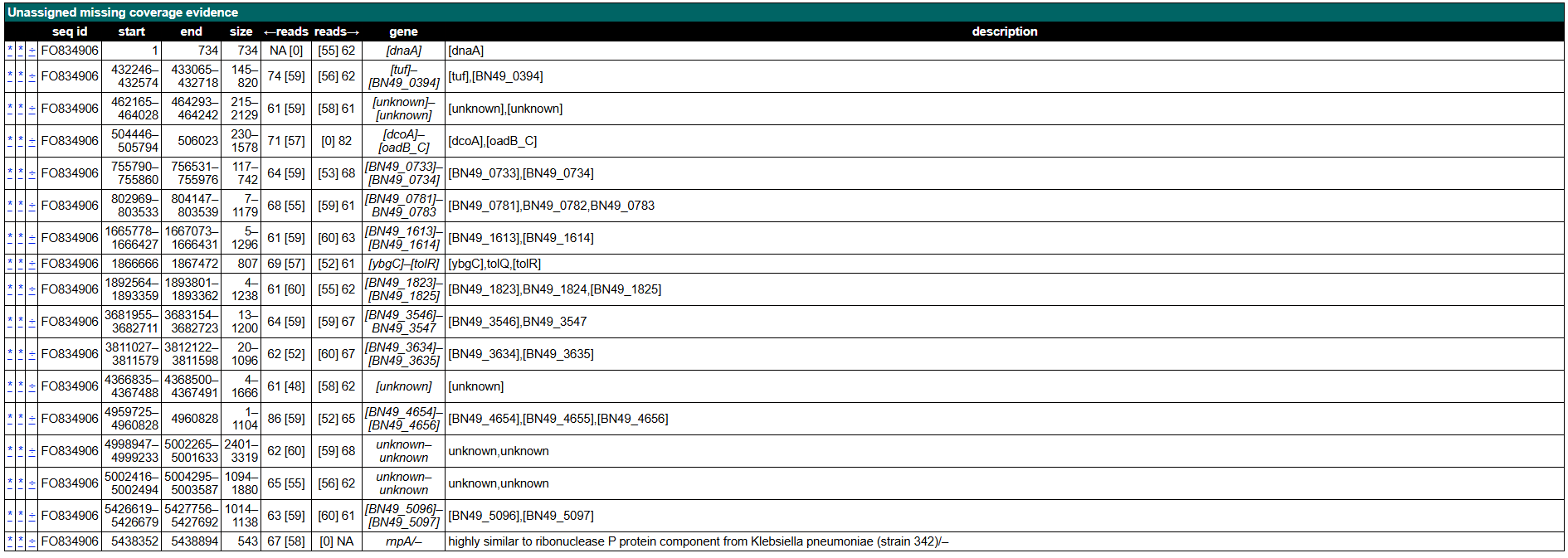

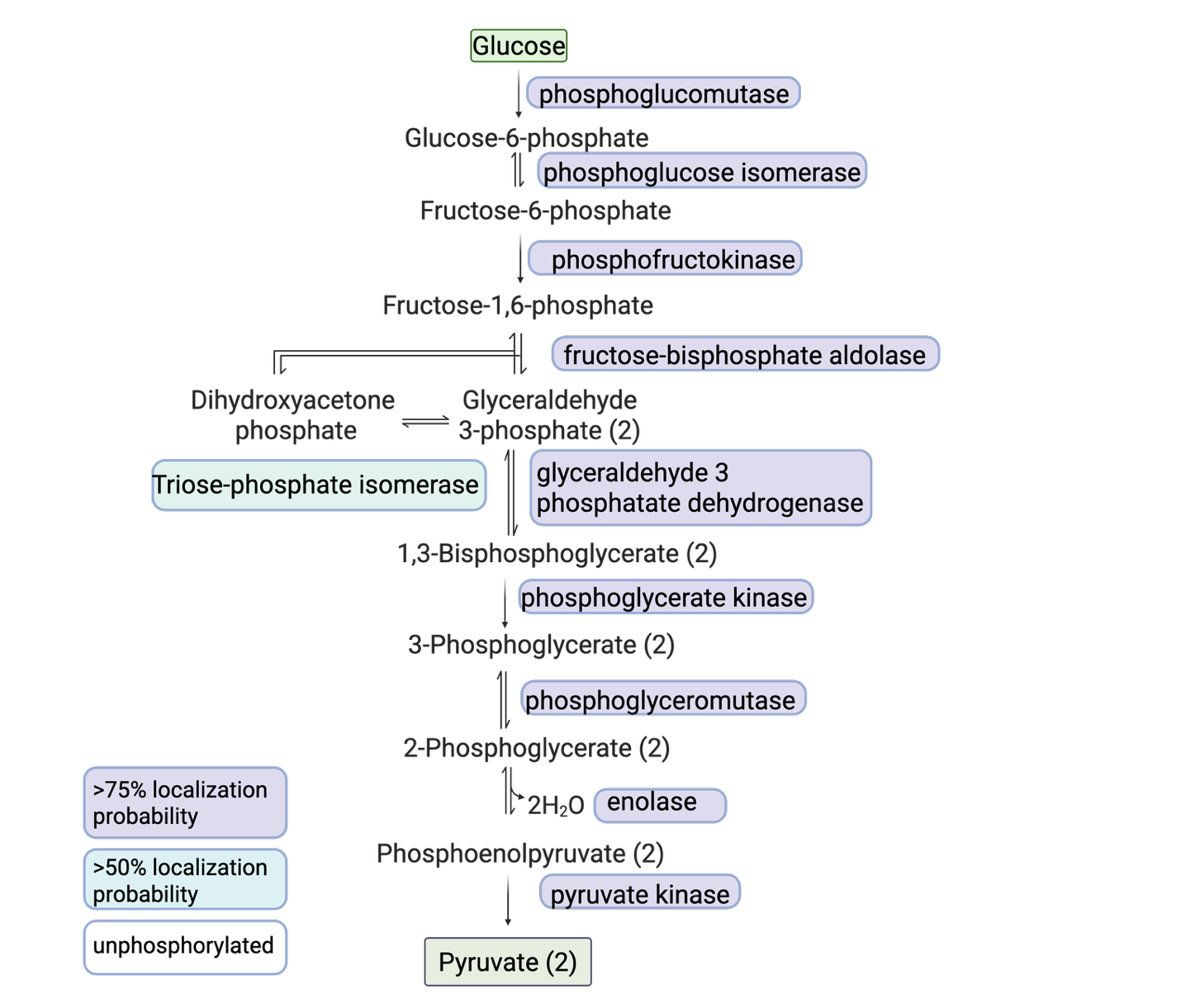

**Figure S1:** Phosphoproteome profiling with identification of known phosphoproteins in glycolysis. Pathway mapping performed using KEGG pathways.

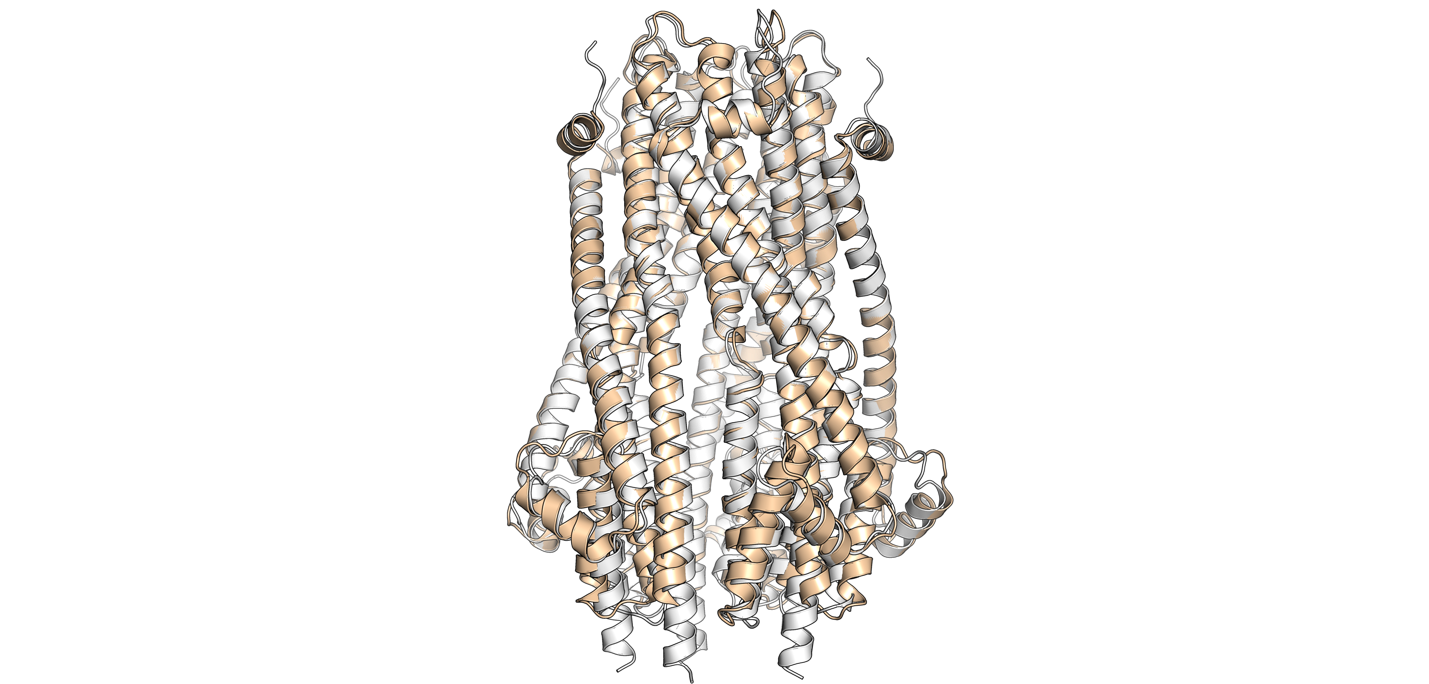

**Figure S2:** *K. pneumoniae* TolQ (A0A0W7ZZF0) superimposed with *A. baumannii* TolQ. The visualization of structures was performed in PyMOL v3.1.5.1.

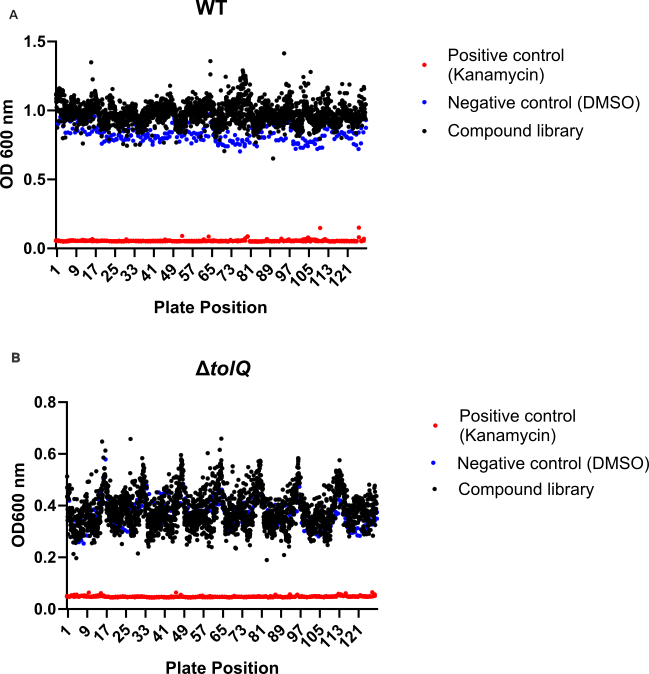

**b**

**a**

**Figure S3:** Optical density (OD_600nm_) of *K. pneumoniae* strains following exposure to the SPARC drug-like curated compound library for 24 h at 37˚C with shaking***.*** (**a**) WT. (**b**) Δ*tolQ.*
